# BCG activation of trained immunity is associated with induction of cross reactive COVID-19 antibodies in a BCG vaccinated population

**DOI:** 10.1101/2024.01.02.573408

**Authors:** Najeeha Talat Iqbal, Kumail Ahmed, Tehniat Sattar, Fatima Aziz, Rabia Hussain

## Abstract

**Background:** During the current COVID-19 pandemic, the rate of morbidity and mortality was considerably lower in BCG vaccinated countries like Pakistan. BCG has been shown to provide cross protection to both disseminated TB as well as non related viral infections in BCG vaccinated children which is consistent with COVID-19 morbidity in the younger age group. Recently, this cross protection was attributed to trained immunity (TI) associated with BCG recall responses in the innate arm of the immune system. Little is known about the longevity of BCG Trained Immunity (TI) beyond early childhood.

**Objective:** To assess the BCG-induced recall responses in healthy individuals by cytokines secreted from the TI network and its potential role in providing cross-protection against COVID-19 and other viral infections.

**Study Design:** In this cross-sectional study, healthy young adults and adolescents (n=20) were recruited from 16–40 years of age, with no prior history of TB treatment, autoimmune, or chronic inflammatory condition.

**Methods:** BCG-induced cytokine responses were assessed using prototypic markers for cells of the TI network {macrophages [M1 (TNFα, IFNγ), M2 (IL10)], NK (IL2), Gamma delta (γδ) T (IL17, IL4)} and SARS CoV2 IgG antibodies against RBD using short-term (12 hrs.) cultures assay.

**Results:** Significant differences were observed in the magnitude of recall responses to BCG with macrophage cytokines showing the highest mean levels of TNFα (9148 pg/ml) followed by IL10 (488 pg/ml) and IFNγ (355 pg/ml). The ratio of unstimulated vs.BCG-stimulated cytokines was 132 fold higher for TNFα, 40 fold for IL10, and 27 fold for IFNγ. Furthermore, SARS-CoV-2 antibodies were also detected in unstimulated plasma which showed cross reactivity with BCG.

**Conclusion:** The presence of cross reactive antibodies to SARS-CoV-2 and the relative ratio of pro-and anti-inflammatory cytokines secreted by activated TI cellular network may play a pivotal role in protection in the early stages of infection as observed during the COVID-19 pandemic in the younger age groups resulting in lower morbidity and mortality.

## 1. Introduction

Pakistan ranks among the top six highest TB burden countries, contributing approximately 56% of the total TB burden globally (1, 2). Pakistan still provides Bacille Calmette-Guerin (BCG) vaccine at birth (>90%) as part of the World Health Organization (WHO) Expanded Program of Immunization (EPI) which was introduced country-wide in 1965. Although BCG protection against pulmonary disease was limited in adults (3–5) but in children, BCG vaccination provides protection against disseminated disease in early childhood (6, 7) as well exhibits non-specific beneficial effects by decreasing mortality in children with respiratory viral illnesses (8), and sepsis (9, 10). For these reasons, BCG vaccine coverage has been continued by WHO in TB endemic countries. BCG vaccination is given at birth, so there are no reliable markers for assessing BCG vaccination status in young adults, as T cells are known to wane over time and most available markers (BCG scar, MT, or PPD) are reflecting T cell memory responses (11). During the COVID-19 pandemic, Pakistan reported relatively low morbidity and mortality rates compared to the rates reported by the Western Countries where no universal vaccination policy exists or BCG has been discontinued (12). It was therefore, speculated that BCG may also provide cross-protection against COVID-19 disease (13–15) and may contribute to decreased morbidity and mortality in Pakistan (12). The rationale for this speculation lies in the fact that BCG is a potent stimulator of the innate arm of the immune system which recognizes conserved antigens (PAMPs) shared across several species (16–18). Although innate immunity was not considered to have memory, recently a concept of TI possessing memory has emerged (19, 20). TI is defined as a bridge between innate and adaptive immune response, predominantly involving innate cells. Recurrent boosting of the innate cellular network in a TB endemic area such as Pakistan may occur due to environmental exposure to TB or non-pathogenic mycobacteria resulting in remodeling and epigenetic changes in monocytes which result in a heightened immune response upon re-infection (21, 22), thus providing protection to other non-related pathogens such as COVID-19 (23). Although there is extensive data available with respect to recall responses in the adaptive or T cell arm of the immune response in BCG vaccinated countries including Pakistan (22), there is little information regarding recall responses in the innate arm (TI) in adults. The cells that form the TI network are cells of the monocyte/macrophages (Mφ) lineage comprising of several subsets (M1, M2a, M2b, and M2c), natural killer (NK) cells, and gamma delta (γδ) T cells derived from the T cell lineage. B cells are also activated by certain antigens to secrete IgG antibodies without T cell help via BCR cross linking and thus form part of the innate network (24, 25). Innate cells are activated by PAMPs via their appropriate Pattern Recognition Receptors (PRRs) (26, 27) (Supplementary Table S1). Cytokines act as growth factors, activators, modulators, and regulators of the immune response, and TI acts as a bridge between the innate and adaptive arms in orchestrating the adaptive arm. We, therefore, investigated BCG induced TI responses by assessing prototypic cytokine secreted by the cells of the innate network. The balance between pro-and anti-inflammatory cytokines may play a key role in disease outcome. Our primary objective was to identify TI activation, using BCG stimulated whole blood assay (WBA) and purified peripheral blood mononuclear cells (PBMCs) cultures (28). Another interesting observation was the presence of BCG cross-reactive IgG antibodies to SARS-CoV-2 antigens. We hypothesize, that the presence of PAMPs like molecules in SARS-CoV-2 may activate the TI network in BCG vaccinated young adults. Both the presence of SARS-CoV-2 cross reacticve IgG antibodies and cytokine secreted by TI network may play a crucial role in determining less severe disease outcome of COVID-19 in young adults as observed during the COVID-19 pandemic in Pakistan and other countries with wide BCG coverage.

## 2. Materials and Methods

### 2.1. Ethics

The research protocol was approved by the Ethical Review Committee of Aga Khan University (ERC protocol 2855-14) Pakistan.

### 2.2. Study Design

The study subjects were recruited from one of the field sites developed by The Department of Pediatrics and Child Health, Aga Khan University, Karachi, during the year 2015-2016. Subjects (n=20) were enrolled after informed consent. The inclusion criteria were healthy adolescents, matched for gender and age (range: 16 - 40 years). Exclusion criteria were, a history of past TB treatment or autoimmune or chronic disease, or steroid therapy for any condition. BCG scar presence was noted, and the Mantoux Test was also conducted (Supplementary Table S2).

### 2.3. Sample Processing

Five ml of blood was used for whole blood assay (WBA) after diluting 1/4 with complete RPMI 1640, while the rest of the blood was used to isolate PBMCs by Ficoll-Isopaque density gradient centrifugation. Media was supplemented with 200mM L-Glutamine, 10X Penicillin-streptomycin mix, 1 M HEPES, and 100 mM Sodium Pyruvate (Sigma Aldrich, St. Louis MO). Cell concentration was adjusted at 2x10^6 cells/ml in 24 well cell culture plates (Corning Incorporated, NY, USA).

### 2.4. BCG stimulation of cell cultures

Freeze-dried Glutamate BCG was obtained from the Serum Institute India. BCG (2-8x10^6 CFU) was reconstituted in 250µl of RPMI to obtain MOI of 1.2 in 60µl of BCG as per Hanekom protocol (29). Freshly reconstituted BCG was added to stimulate the peripheral blood mononuclear cells (PBMCs) (2x10^6) and whole blood (WB) (1/4 diluted) and incubated for 12 hrs. Supernatants were harvested and stored at -80 LC until further use. Additionally, WB and PBMCs were stimulated using 10µg/ml of Lipopolysaccharide (LPS) (Sigma Aldrich, St. Louis MO).

### 2.5. Assessment of cytokine responses in BCG induced cell cultures

BCG stimulated and un-stimulated cell cultures were assessed for secreted cytokines post 12 hrs. incubation. Cytokines were analyzed on Bio-Rad Luminex 200 platform. The bead mixed panel includes IL2, IL4, IL10, IL17A, IFN_γ_, and TNF_α_. Data were acquired using Bio-Plex Manager Software version 6.1 (Bio-Rad).

### 2.6. Gene Expression analysis for BCG stimulated cells using real Time PCR

#### 2.6.1. RNA Extraction

RNA extraction was carried out after RBC lysis from WBA and PBMCs culture using TRIzol reagent (Ambion, life technologies, Carlsbad CA, USA). Finally, the RNA pellet was re-suspended in 20µl of DNase RNase-free water (Gibco, Life Technologies, NY, USA). RNA purity and quantity were checked by Nanodrop 2000c.

#### 2.6.2. cDNA synthesis

cDNA was synthesized using the Bio-Rad Iscript buffer (Bio-Rad Lab. Hercules, CA) as per the manufacturer’s instruction. The protocol was run on the CFX 96 Bio-Rad (Hercules, CA) platform.

#### 2.6.3. Real-time quantitative PCR

The oligonucleotide primers were designed from IDT for Granzyme (GZMA), Butyrophilin (BTN3A2), Tumor Necrosis Factor (TNF_α_), and 36B4 (as housekeeping) genes. qPCR was performed using Bio-Rad SYBR Green (Bio-Rad Lab. Hercules, CA) as per the manufacturer’s instructions. Briefly, cDNA was used as a template with a 1:10 ratio of cDNA and SYBR green in a total of 25µl of reaction and read at Bio-Rad CFX 96 platform. All results of the test gene were normalized with a housekeeping gene (36B4 gene). ddCt was used to express gene expression after normalizing with the housekeeping gene. Primers used to analyze different genes are shown in (Supplementary Table S3).

### 2.7. Assessment of IgG anti-RBD (SARS-CoV-2) in Plasma using Elisa assay

Immulon 4 ELISA plates were coated with receptor binding domain (RBD) antigen (Institute of Protein Design-IPD, The University of Washington Seattle USA) (2 µg/ml) in 1X PBS as coating buffer and incubated overnight at 4 °C. Plates were washed with 1X PBS-0.1% Tween20 (PBST) and blocked with 2% bovine serum albumin (BSA). Plasma samples were diluted 1:500 in 1X PBST-2% BSA and added to their respective wells. Plates were incubated for 1.5 hours at room temperature with shaking. Plates were subsequently washed with 1X PBST during each step. Goat Anti-human-IgG (H+L) secondary antibody conjugated with horse-reddish peroxidase was added at 1:30000 dilution and plates were incubated for 1 hour at room temperature while shaking. Plates were finally developed using o-phenylenediamine (OPD) tablets mixed with sodium perborate buffer (Sigma-Aldrich). All plates were read on a MultiSkan Sky ELISA plate reader (ThermoFisher Scientific MA USA) at 492 nm. A dose response curve was developed using a high titer of positive IgG antibodies pool from COVID-19 patient’s samples. The positive pool was serially diluted from 1:1000 to 1:128000 and an unknown concentration of IgG was read using the standard curve. R^2^ was 0.9954 and the negative pool has less than 0.3 O.D. Samples were considered positive for IgG if they had higher Mean +3SD from the negative pool.

### 2.8. Data analysis

The data were analyzed using a non-parametric test between the subjects using the Mann-Whitney U test and Wilcoxon Sign Rank test for within subjects. Spearman rank correlation analyses were applied for a significant correlation between cytokines. Principal component analyses using factor analyses to cluster cytokines into different groups. IBM Corp. Released 2012. IBM SPSS Statistics for Windows, Version 21.0. was used for data entry and analyses. R Studio Package, Version 4.1.2, (www.r-project.org) was used for data visualization using heat maps and dot plots. Cytokine data was reported after the deduction of background signal from un-stimulated controls. Signal to Noise ratio S/N was calculated by dividing the mean of stimulated cytokine level by mean of spontaneous or background secretions. Cytokine or gene expression data was reported after normalization of the test gene by the housekeeping gene for stimulated and un-stimulated controls, the results are expressed as an arbitrary unit (AU) of the expressed gene relative to the expression of negative or un-stimulated control using the ddCt method.

## 3. Results

### 3.1. Characteristics of the study group

COVID-19 pre-pandemic healthy adults (n=20) were recruited between 2015 to 2016. Gender distribution was similar, and the median age was 27 years. BCG scar and MT frequency were 40% and 10% respectively. This is not surprising as T cell responses wane over time. MT positivity was much lower (10%) than BCG scar probably because of the low sensitivity of MT (Table 1).

**Table 1:** Characteristics of study participants.

| Study Subjects | n=20 |
| --- | --- |
| Male / Female | 10 / 10 |
| TB Contact History | 2 / 20 |
| BCG Scar | 8 / 20 |
| MT positivity (> 5.0 mm) | 1 / 20 |
| Age: Median; Range (Years) | 27; (16 – 40) |
| Cell count (PBMCs per ml) mean ± SE | 190.023 ± 2.23 |

Note: All participants were enrolled pre-COVID-19 pandemic

Since there is >90% BCG vaccination coverage at birth in Pakistan, neither BCG scar nor MT positivity was reflective of BCG vaccinated status. Since there was also no difference in cytokine responses between BCG vaccinated and non-vaccinated subjects (Supplementary Fig. S1). We, therefore, carried out the TI analyses in the un-stratified study.

### 3.2. Cytokine secretion in un-stimulated and BCG stimulated whole blood assay (WBA) and PBMCs cultures post 12-hour incubation

BCG induced recall responses were observed in all cells of the TI network. Fig. 1 shows the complete dynamics of cytokine secretion for both assays. Spontaneous cytokine activation is shown as a line over the shaded area representing BCG stimulation. All cytokines showed recall responses but there were several log differences in cytokines secreted by cells of monocyte (TNF_α_, IFN_γ_, IL10) and cells of the T cell lineage (IL2, IL4, IL17). When BCG stimulated cytokines were compared in the two assays, TNF_α_, IFN_γ_, IL2, and IL4 were up-regulated in the WBA culture compared to the PBMCs culture. On the contrary, IL17 (p = 0.003) and IL10 (p *=* 0.013) showed significantly higher secretion in PBMCs (Fig. 1-shaded area).

**Fig. 1.**
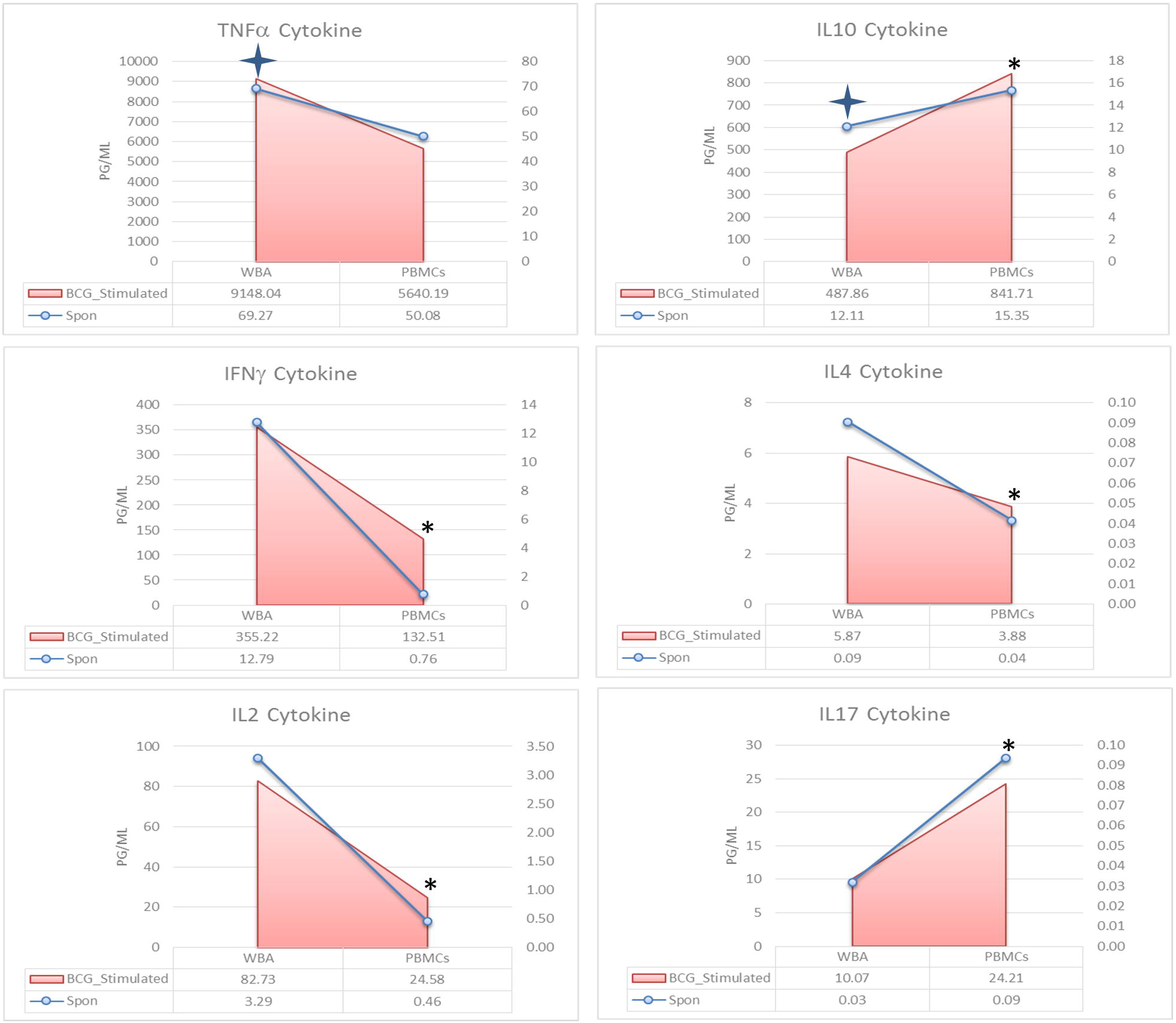
Cytokine secretion in spontaneous and Bacille Calmette-Guerin (BCG) stimulated Whole Blood Assay (WBA) and peripheral blood mononuclear cells (PBMCs) short-term culture supernatants. The shaded area compares secretion of WBA and PBMCs (n=20) as the mean level of individual cytokine in BCG stimulated cultures (12 hrs.) shown on the primary axis (left side). The line graph compares the spontaneous secretion of the cytokine to be read on the secondary axis (right side). The “4-sided star” indicates a significant *p*-value for spontaneous cytokine secretion, whereas “asterisks” shows the difference in BCG stimulated cytokines in WBA and PBMCs. Wilcoxon Sign Rank test was applied for the significant difference in the cytokine level. *p* < 0.05 was considered significant.

When WBA and PBMCs cultures were compared for spontaneous cytokine secretion (Supplementary Table S4), significant differences were noted for macrophage cytokines with higher responses in WBA for TNF_α_ (MWU; *p* = 0.009) and significantly lower IL10 (MWU; *p*= 0.017) in WBA compared to PBMCs. Because of differential spontaneous secretion in the two assays, we analyzed net responses which probably reflect true recall responses. This analysis provided a different set of answers. The highest concentrations (*p*g/ml) were again observed with macrophage secreted cytokines but with more clear differences in the mean level of cytokines in two assays (WBA vs PBMCs: TNF_α_, 9148 vs 5640; IFN_γ_, 355 vs 132; IL10 488 vs 842; IL2, 83 vs 24; IL4, 6 vs 4; IL17, 10 vs 24) (Supplementary Fig. S2).

We also looked at recall responses in terms of the ratio of increase post stimulation (Signal/Noise) in the two assays. Four of six cytokines showed comparable S/N ratio for WBA and PBMCs (IL2, IL4, IL-10 and IL17) while differences in S/N ratio were higher for TNF_α_ in WBA (132 –fold) compared to PBMCs (113-fold). IFN_γ_ on the other hand showed lower concentrations in WBA compared to PBMC with a higher ratio for PBMC (WBA; 27 v s PBMCs 132) (Supplementary Table S5). So, both the magnitude and ratio of recall response are higher for macrophage secreted cytokines in WBA compared to PBMCs indicating that WBA may provide a better window for assessing recall responses.

### 3.3. Effect of BCG scar on BCG stimulated cytokine secretion

We also compared cytokine responses in subjects stratified by BCG scar positivity. Although the group sizes are relatively small, no significant differences were noted in cytokine secretion in WBA for the two groups (Supplementary Fig. S1). This is not surprising as the presence of BCG scar is related to adaptive T cell responses and not to innate immunity.

### 3.4. Effect of BCG scar on gene expression in stimulated PBMCs

We also carried out prototypic gene expression analysis associated with TNF_α_ (macrophages), Granzyme A (GZMA; NK cells), and Butyrophilin (BTN3A2; gamma delta (γδ) T cells) in PBMCs (Supplementary Table S3). TNF_α_ was the only gene which showed differential upregulation in the BCG scar negative subjects compared to the BCG scar positive subjects (p = 0.038). The implications of up regulated TNF_α_ expression in BCG scar negative donors and its association with BCG vaccination status is unclear. There was no difference in gene expression for natural killer (NK) cells and gamma-delta (γδ) T cells (Fig. 2).

**Fig. 2.**
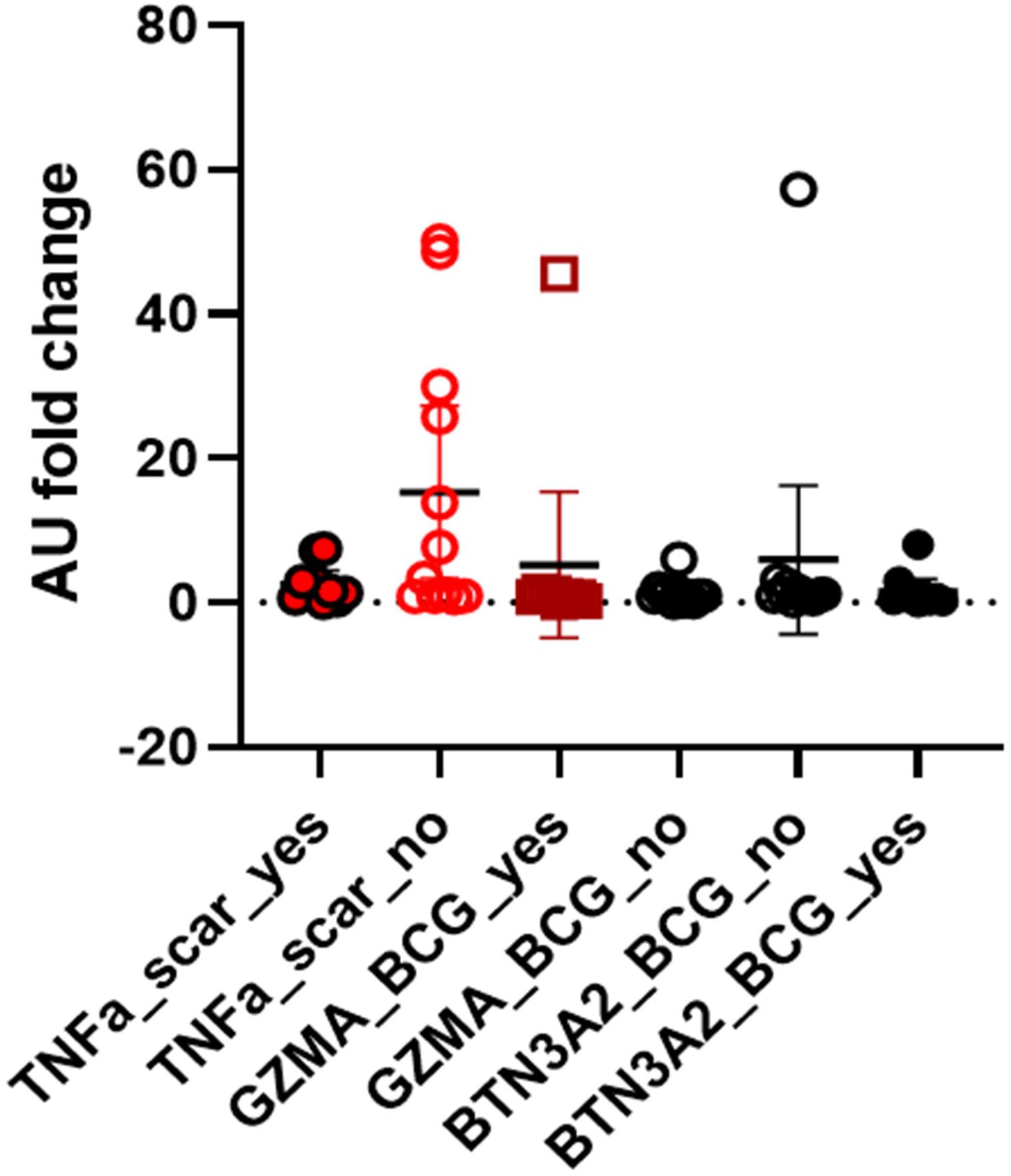
Expression of innate and adaptive genes in response to Bacille Calmette-Guerin (BCG). The scatter plot shows the fold changes in the expression of Tumor Necrosis Factor (TNF_α_), Granzyme A (GZMA), and Butyrophilin (BTN3A2) genes in response to live BCG stimulation of PBMCs. The fold changes are shown as relative expression compared to non-stimulated media control (10% FBS) using 2x10^6 cells between 12-24 hrs. Wilcoxon Sign Rank test were applied for the comparison of Scar positive and negative subjects.

#### 3.4.1. Principal component analysis (PCA)

We next addressed the issue of the relationship between different cytokines in the two assays. Principal Component Analysis (PCA) was carried out to determine cytokine clustering patterns in WBA and PBMCs (Fig. 3). PCA separated data into two components PC1 which contained 5/6 cytokines and PC2 with IL10 in WBA cultures. Cytokines secreted from PBMCs are clustered as a single component. This is interesting as PC1 clusters all the pro-inflammatory cytokines which are derived predominantly from M1 macrophages, while IL10 which is an anti-inflammatory cytokine is secreted not only from M2 macrophages but several other cell sources of the TI network. Therefore, PC analysis confirms our previous contention that TNF_α_ and IL10 are from different cell sources.

**Fig. 3.**
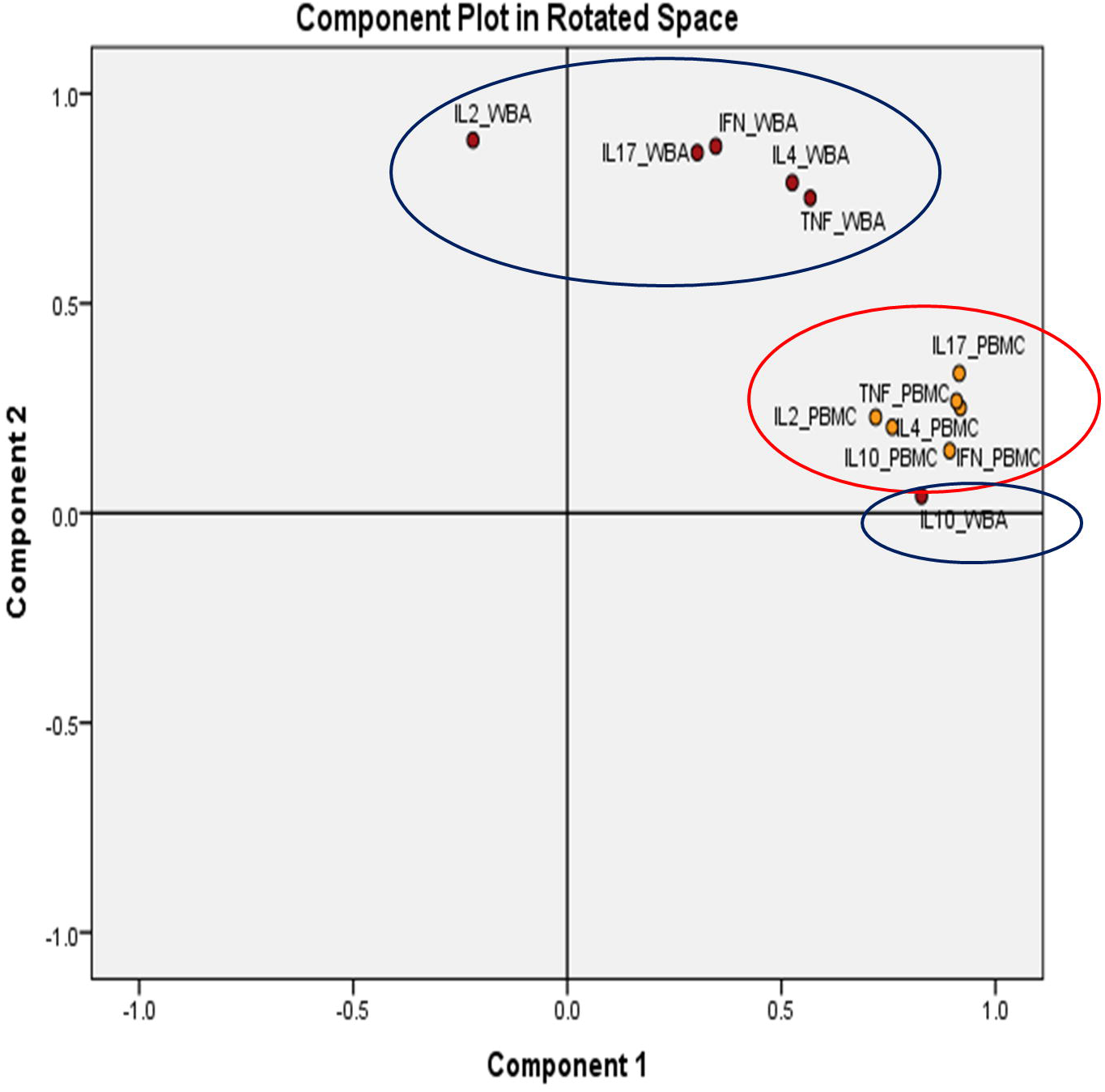
Assay dependent clustering pattern of BCG stimulated cytokines. The orange dots indicate cytokines analyzed through PBMCs while the red dot indicates cytokines analyzed through WBA. The PCA separated data into two major components. All variables explained 80% of the variability in the model.

#### 3.4.2. Correlation analysis of secreted cytokines in WBA and PBMCs cultures

To further understand the relationship between secreted cytokines in WBA and PBMC, we carried out correlation analyses. (Fig. 4). Very similar patterns were observed with both WBA (4A) and PBMCs (4B).

**Fig. 4.**
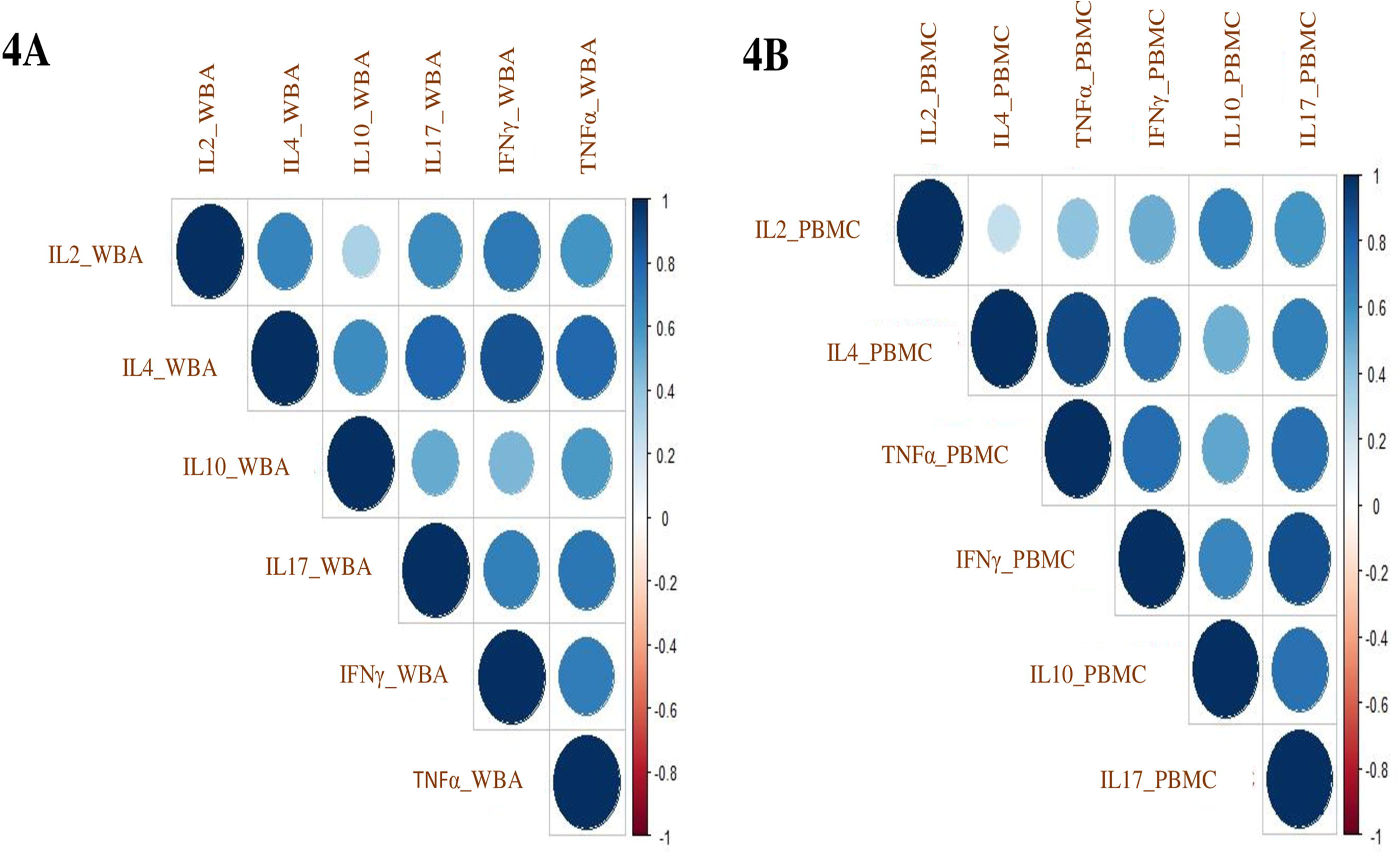
Cytokine correlation index in WBA and PBMCs cultures. Corrplot showing a correlation coefficient of cytokines using the intensity of colors at the scale of -1.0 to +1.0 in WBA (**A**) and PBMCs assay (**B**).

The correlation indices are shown in Supplementary Tables S6 and S7. All cytokines showed a highly significant positive correlation (>0.5), except for IL10 in WBA (highest, r=0.515 with IL4; lowest r=0.277 with IL2), and PBMCs (highest, r=0.681 with IL17; lowest r=0.455 with IFN_γ_). This is consistent with what we observed in PCA analysis with IL10 grouping differently than all other cytokines. Differences in PC compartments and differential correlation in WBA and PBMCs culture suggest that WBA culture may be a more appropriate window for investigating TI responses as it is better reflective of in vivo situations due to minimal processing.

### 3.5. Comparison of cytokine secretion in WBA cell culture in response to BCG and LPS stimulation

Lipopolysaccharide (LPS) is a potent mitogen and a strong stimulator of macrophages. We next compared BCG and LPS stimulated cytokine secretion to identify if there was any commonality in antigen recognition between those antigens. The concordance of antigen recognition with BCG and LPS is shown in Fig. 5. Responses with LPS were comparable for TNF_α_, and IFN_γ_, while IL17, IL2, and IL4 showed a much lower overall magnitude of responses with LPS antigen. IL10 was higher in LPS stimulated samples compared to BCG. Interestingly, BCG was a better stimulus compared to LPS with IL17, IL2, and IL4.

**Fig. 5.**
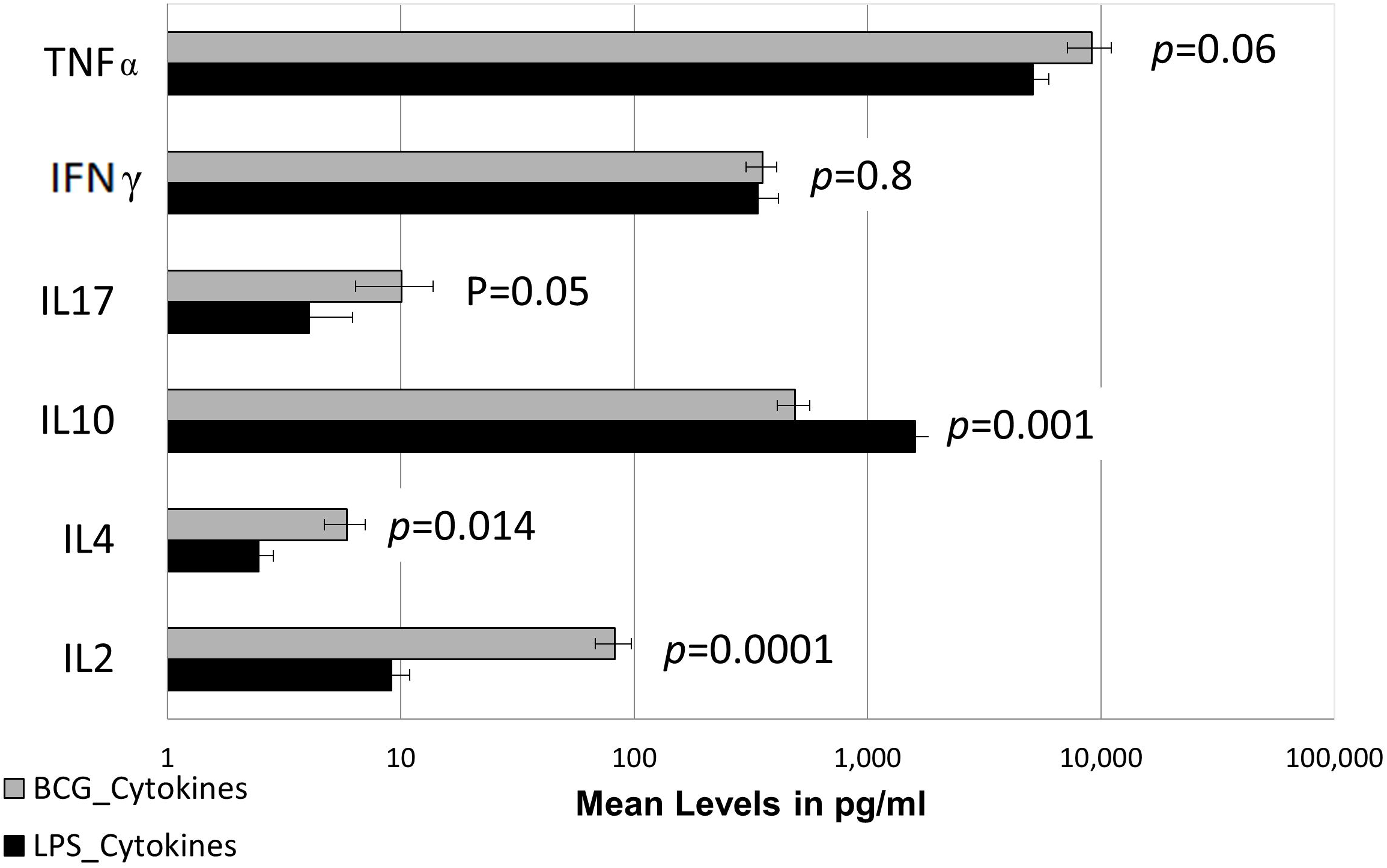
Comparative analysis of Bacille Calmette-Guerin (BCG) and Lipopolysaccharide (LPS) stimulated cytokine secretion in Whole Blood Assay (WBA) cell culture. Comparison of cytokine secretion in response to BCG and LPS stimulation in WBA cell cultures at 12 and 48 hrs. respectively (n=20). Horizontal bars indicate mean levels with standard error bars around the mean. Student t-tests were carried out to assess the significance of differences between BCG and LPS.

As expected, the highest correlation between LPS and BCG was observed with TNF_α_ (SR rho= 0.67; *p =* 0.002) followed by IL4 (SR rho = 0.611; *p* = 0.005), IL2 (SR rho = 0.562; *p* = 0.015) and IFN_γ_ (SR rho = 0.464; *p* = 0.045). Again, no correlation was observed with IL10 (SR rho= -0.033; *p* = 0.896) and IL17 (SR rho = 0.273; *p* = 0.258) (Supplementary Table S8). These results strongly suggest that all induced cytokines are recognizing PAMPs like molecules and are derived from cells of the TI network.

### 3.6. Assessment of IgG anti-RBD antibodies in the plasma of healthy donors

Since it has been reported that BCG provides non-specific immunity against other respiratory infections, we investigated if cross-reactive IgG against Receptor Binding Domain (RBD) were present. A calibration curve was generated with a pool of COVID-19 positive sera (Supplementary Fig. S3) with the endpoint titer considered as 1 unit/ml.

All healthy donors showed considerable levels of SARS-CoV-2 RBD antibodies (Fig. 6). The binding of antibodies to RBD was inhibited in the presence of BCG suggesting recognition of cross-reactive PAMPs like molecules in RBD.

**Fig. 6.**
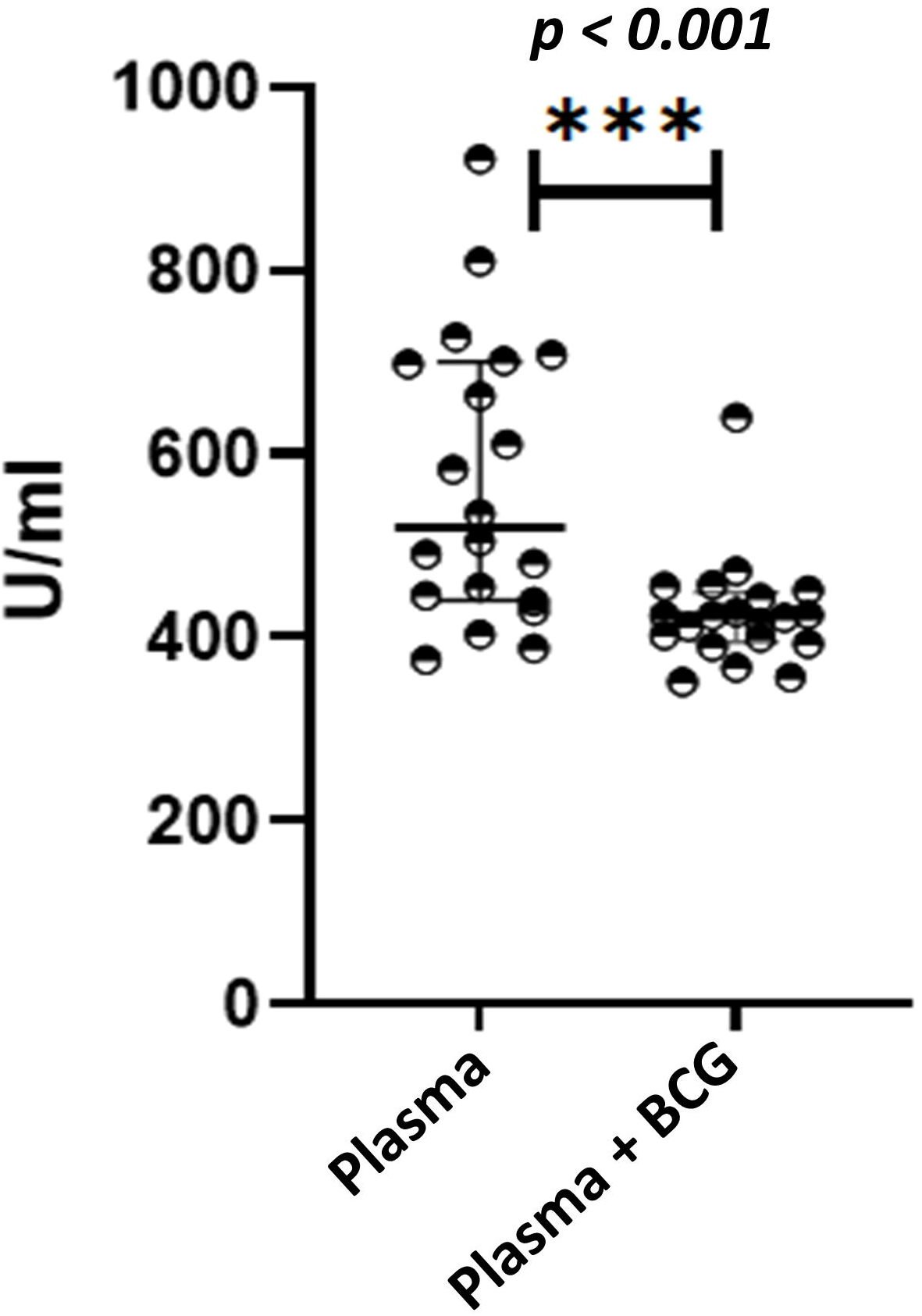
Assessment of cross-reactive IgG antibodies to COVID-19 Receptor-Binding Domain (RBD) Results are expressed as units /ml. Antibody assessment was carried out in plasma and BCG stimulated supernatants. All samples were collected before the COVID-19 pandemic. Mann-Whitney U tests were carried out to determine the significance of the difference.

### 3.7. Relationship of IgG with Cytokines

The correlation plot (Fig. 7) shows the relationship of IgG anti-RBD with BCG and LPS-driven cytokines in WBA. There was no relationship between IgG with either IL10 or IL4 which are considered growth factors for B cells. However, IL17 (r = -0.791; *p =* 0.001) and IFN_γ_ (r = -0.717; *p* = 0.005) showed a strong negative relationship with IgG anti-RBD. We believe that IgG recognition is not only T independent but recognizes different cross-reactive epitopes such as carbohydrate moieties and therefore shows a totally independent response compared to BCG and LPS induced cytokines.

**Fig. F7.**
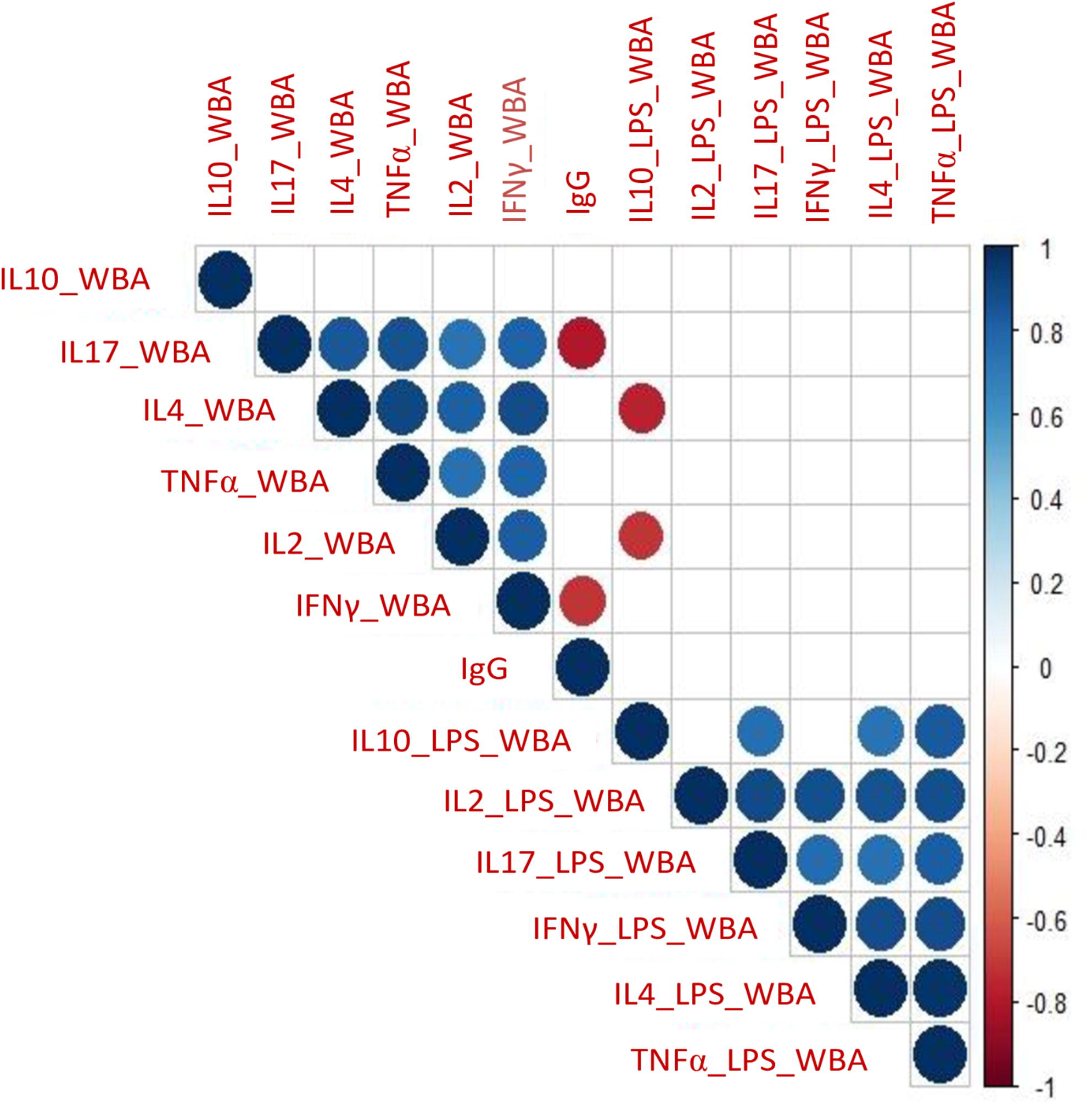
Relationship of IgG anti-RBD with BCG and LPS driven cytokines in Whole Blood Assay (WBA) Correlation plot of cytokines derived from WBA (BCG and LPS) and IgG. Corrplot showing a correlation coefficient of cytokines using the intensity of colors at the scale of -1.0 to +1.0 in WBA. All significant correlation at the level of *p* = 0.01 is shown on the plot as circles and *p* > 0.01 are shown as blank.

## 4. Discussion

In this study, we show BCG-induced activation of the TI cellular network across the board in healthy adult donors in a BCG vaccinated area. This study is unique in the sense that case recruitment was conducted between 2015-2016 before the onset of COVID-19 pandemic. Secondly, while most previous studies have focussed on BCG vaccinated paediatric group, we have extended the age range to young adults to address the issue of longevity of BCG recall responses in TI network. This study therefore provides information related to BCG related longevity of TI recall responses in young adults pre COVID-19 pandemic in a TB endemic country, and is not possible to obtain post COVID-19. The importance of TI is the early activation (minutes to hours) of encountering a danger signal and can play a crucial role in stopping the infection in its track at the initial stages (30, 31).

One limitation of this study was the lack of an objective marker to determine BCG vaccination status in adults. Neither BCG scar (40% positive) nor MT (10%) could provide this information as these markers are related to T cells which wanes over time (11). Interestingly, TNF_α_ gene expression was the only biomarker which showed differential expression in BCG Scar negative and Scar positive donors. Since kinetics of cytokine gene expression is different for different cytokines more detailed analyses are required to understand the implications of this finding in relation to BCG scar and vaccination status. However, with a >90 percent coverage of BCG vaccination at birth in Pakistan, we feel confident that the majority of our donors were vaccinated, and we are observing BCG recall responses for different cytokines in these healthy donors. This is further supported by similar cytokine secretion in BCG scar negative and BCG scar positive subjects (Supplementary Fig. S1).

Cytokine activation was most marked with the classical (M1) and alternatively activated (M2) macrophages as demonstrated by the pro-inflammatory cytokines TNF_α_ and IFN_γ_ (M1) and immunosuppressive cytokine IL10 (M2) (32). Previously we have shown that IFN_γ_ / IL10 plays a critical role in TB disease progression (33). The ratio observed in the current study is consistent with a protective ratio. IFN_γ_ is also contributed by NK cells and NK activation may play a critical role in viral uptake and killing, thus reducing the burden of viral load at the initial stages of infection. In addition, there was also BCG activation of γδ T cells (IL4 and IL17) but lower compared to Macrophage activation (TNF_α_ and IL10). However, limited activation was observed for NK cells (IL2) (16, 34). The redundancy in cytokine secretion from different sources makes it difficult to evaluate individual contribution of cell subtypes. We addressed this issue by analyzing clustering patterns using Principal component analysis (PCA). The fact that these cytokines are from distinct sources was confirmed by the distinct clustering patterns observed with TNF_α_ / IFN_γ_ clustering as one component (PC1) and IL10 as a second component (PC2) in WB. Surprisingly, PBMCs showed only one cluster. Purification of PBMCs has been shown to result in a loss of adherent cells. (35, 36). The decrease in IFN_γ_ and increase in IL10 in PBMC suggests a selective loss of M1 cells. This observation is also consistent with the known cross regulation of IFN_γ_ and IL10. Again, these observations reinforce the point that WBA may be more reflective of in vivo situations.

Kleinnijenhuis et al., 2014 (37) were the first to describe TI in BCG vaccinated healthy donors to unrelated pathogens. The main difference between innate and adaptive immunity is that cells of the TI recognize antigens shared across pathogens and do not require a second signal as is the case with adaptive immunity. Comparison of BCG-induced responses with LPS which is present in bacterial cell walls and is similar to the lipid coat of mycobacteria showed similar responses for the two antigen confirming strong recognition of conserved epitopes by innate network. As expected, the two cytokines associated with M1 macrophages (TNF_α_ and IFN_γ_) showed a high level of correlation with BCG and LPS-induced TNF_α_, supporting the recognition of PAMPs like moieties in both molecules (38). We have only analyzed prototypic cytokines, but there are other cytokines such as IL1 and IL6 which are also secreted by these macrophages and may also play a role in regulating and orchestrating the adaptive arm. However, the extended set of cytokines was not within the scope of our study.

Up-regulation of the pro-and anti-inflammatory arm of the innate system may serve a crucial role in creating an appropriate balance and dictating the evolution of the disease process. BCG recall responses assessed as a ratio of TNF_α_/IL10 was 3.3 and 2.0 in WBA and PBMCs respectively (Supplementary Table S5). A balanced activation of pro-and anti-inflammatory cytokines at the initial stages of the disease may play a key role in determining disease outcomes to a variety of cross-reactive pathogens including SARS-CoV-2 infections (39). In this context, we have also observed BCG cross-reactive antibodies to SARS-CoV-2 RBD antigen. None of the cytokines showed any relationship with IgG anti-RBD although both IL4 and IL10 are B cell growth factors in the adaptive arm of the immune response (40). Therefore, this is most likely memory responses of long term T independent memory B cells which are activated directly by cross reactive B cell epitopes (41). Such cross-reactive IgG anti-SARS-CoV-2 RBD antibodies may be able to restrict viral replication by forming immune complexes and facilitating entry into macrophages and NK cells at the initial stages. Dissecting the nature of IgG subtypes may shed further light into protective vs pathogenic antibodies. BCG is a cross reactive mycobacteria and BCG vaccination was given with the assumption that the cross reactive response will be sufficient for protection against M. tuberculosis. BCG however, is not effective against pulmonary tuberculosis, but was shown to provide protection against disseminated and miliary disease in children and was therefore continued in TB endemic countries. Protection against disseminated tuberculosis is a function of the innate arm and may be responsible for providing cross protection to other pathogens at entry. This may be one explanation for the protection against non-related pathogens in BCG vaccinated children. There are widely different policies for BCG vaccination in different countries and therefore may result in variable immune responses and protection. Although there is wide coverage globally with BCG (2), the main differences may lie in the community exposure to not only TB but to nonrelated pathogens in endemic countries which may play a key role in re-stimulation of the innate arm of the immune system in healthy donors.

The strongest immune response is shown for Heat Shock Proteins (HSPs) which are conserved across species (42). HSP activates cells of the trained immunity network as well as T cells of the adaptive arm. It is not surprising that immune responses to HSPs have been shown to correlate with a favorable outcome of COVID-19 disease (43).

Further work needs to be done to evaluate the key cells and epitopes which are driving the cytokine TI network particularly as it relates to macrophage subsets and B cells and its implications in cross protection to other non-related pathogens.

## 5. Conclusion

The presecnce of cross reactive antibodies to SARS-CoV-2 and the relative ratio of pro-and anti-inflammatory cytokines secreted by activated TI cellular network may play a pivotal role in protection in the early stages of infection such as observed during the COVID-19 pandemic in the younger age groups resulting in lower morbidity and mortality.

## Acknowledgement

Funding support from Aga Khan University Research Council, Seed Money Grant. Infectious Disease Research lab for logistics and partial funding support for cytokine assays. BCG Japan for provision of BCG vaccine. Professor Van Voorhis for provision of RBD antigen from Institute of Protein Design UW-Seattle. Dr. Anita K M Zaidi, Dr. Farheen Quadri, and Dr. Ali Faisal Saleem for field site support for sample collection. We acknowledge the valuable contribution of Dr. Zehra Jamil in the analysis and discussion part of this paper.

## References

1. WHO. Global Tuberculosis Report 2019-Annexure 2. Country profiles for 30 High TB Burden Countries. 2019 17 Oct 2019.

2. WHO. Global tuberculosis report 2021. Geneva: World Health Organization 2021 2021.

3. Fine PE, Carneiro IA, Milstien JB, Clements CJ, Organization WH. Issues relating to the use of BCG in immunization programmes: a discussion document. Geneva: World Health Organization; 1999.

4. Hasan Z, Irfan M, Khan JA, Jahangir SK, Haris M, Ashraf M, et al. BCG vaccination is associated with decreased severity of tuberculosis in Pakistan. International journal of mycobacteriology. 2012;1(4):201–6.

5. de Gijsel D, von Reyn CF. A breath of fresh air: BCG prevents adult pulmonary tuberculosis. International Journal of Infectious Diseases. 2019;80:S6–S8.

6. Brewer TF. Preventing tuberculosis with bacillus Calmette-Guerin vaccine: a meta-analysis of the literature. Clinical Infectious Diseases. 2000;31(Supplement_3):S64–S7.

7. Aaby P, Roth A, Ravn H, Napirna BM, Rodrigues A, Lisse IM, et al. Randomized trial of BCG vaccination at birth to low-birth-weight children: beneficial nonspecific effects in the neonatal period? Journal of Infectious Diseases. 2011;204(2):245–52.

8. Stensballe LG, Nante E, Jensen IP, Kofoed P-E, Poulsen A, Jensen H, et al. Acute lower respiratory tract infections and respiratory syncytial virus in infants in Guinea-Bissau: a beneficial effect of BCG vaccination for girls: community based case–control study. Vaccine. 2005;23(10):1251–7.

9. Hack CE, De Groot ER, Felt-Bersma R, Nuijens JH, Strack Van Schijndel R, Eerenberg-Belmer A, et al. Increased plasma levels of interleukin-6 in sepsis [see comments]. 1989.

10. Jason J, Archibald LK, Nwanyanwu OC, Kazembe PN, Chatt JA, Norton E, et al. Clinical and immune impact of Mycobacterium bovis BCG vaccination scarring. Infection and immunity. 2002;70(11):6188–95.

11. Whittaker E, Nicol MP, Zar HJ, Tena-Coki NG, Kampmann B. Age-related waning of immune responses to BCG in healthy children supports the need for a booster dose of BCG in TB endemic countries. Scientific reports. 2018;8(1):1–10.

12. Wickramasinghe D, Wickramasinghe N, Kamburugamuwa SA, Arambepola C, Samarasekera DN. Correlation between immunity from BCG and the morbidity and mortality of COVID-19. Tropical Diseases, Travel Medicine and Vaccines. 2020;6(1):17.

13. O’Neill LA, Netea MG. BCG-induced trained immunity: can it offer protection against COVID-19? Nature Reviews Immunology. 2020;20(6):335–7.

14. Mohapatra PR, Mishra B, Behera B. BCG vaccination induced protection from COVID-19. indian journal of tuberculosis. 2021;68(1):119–24.

15. Messina NL, Germano S, McElroy R, Rudraraju R, Bonnici R, Pittet LF, et al. OffLJtarget effects of bacillus Calmette–Guérin vaccination on immune responses to SARSLJCoVLJ2: implications for protection against severe COVIDLJ19. Clinical & translational immunology. 2022;11(4):e1387.

16. Kleinnijenhuis J, Quintin J, Preijers F, Benn CS, Joosten LA, Jacobs C, et al. Long-lasting effects of BCG vaccination on both heterologous Th1/Th17 responses and innate trained immunity. Journal of innate immunity. 2014;6(2):152–8.

17. Arts RJ, Moorlag SJ, Novakovic B, Li Y, Wang S-Y, Oosting M, et al. BCG vaccination protects against experimental viral infection in humans through the induction of cytokines associated with trained immunity. Cell host & microbe. 2018;23(1):89–100. e5.

18. Aaby P, Benn CS. Saving lives by training innate immunity with bacille Calmette-Guerin vaccine. Proceedings of the National Academy of Sciences. 2012;109(43):17317–8.

19. Cassone A. The Case for an Expanded Concept of Trained Immunity. mBio. 2018;9(3):e00570–18.

20. Marcenaro E, Ferranti B, Falco M, Moretta L, Moretta A. Human NK cells directly recognize Mycobacterium bovis via TLR2 and acquire the ability to kill monocyte-derived DC. International immunology. 2008;20(9):1155–67.

21. Hussain R, Toossi Z, Hasan R, Jamil B, Dawood G, Ellner J. Immune response profile in patients with active tuberculosis in a BCG vaccinated area. The Southeast Asian Journal of Tropical Medicine and Public Health. 1997;28(4):764–73.

22. Iqbal NT, Hussain R. Non-specific immunity of BCG vaccine: a perspective of BCG immunotherapy. Trials in Vaccinology. 2014;3:143–9.

23. Tran K, Divangahi M. BCG vaccination provides cross-protection against influenza infection through trained adaptive immunity. The Journal of Immunology. 2021;206(1_Supplement):110.18–18.

24. Alberts B JA, Lewis J, et al. Molecular Biology of the Cell: Helper T cells and Lymphocyte Activation. New York : Garland Sciences (2022).

25. Hess C, Winkler A, Lorenz AK, Holecska V, Blanchard V, Eiglmeier S, et al. T cell– independent B cell activation induces immunosuppressive sialylated IgG antibodies. The Journal of Clinical Investigation. 2013;123(9):3788–96.

26. van der Meer JW, Joosten LA, Riksen N, Netea MG. Trained immunity: a smart way to enhance innate immune defence. Molecular immunology. 2015;68(1):40–4.

27. Suliman S, Geldenhuys H, Johnson JL, Hughes JE, Smit E, Murphy M, et al. Bacillus Calmette–Guerin (BCG) revaccination of adults with latent Mycobacterium tuberculosis infection induces long-lived BCG-reactive NK cell responses. The Journal of Immunology. 2016;197(4):1100–10.

28. Hussain R, Jamil S, Dockrell H, Chiang T, Hasan R. Detection of high titres of Toxoplasma gondii antibodies in sera of patients with leprosy in Pakistan. Transactions of the Royal Society of Tropical Medicine and Hygiene. 1992;86(3):259–62.

29. Hanekom WA, Hughes J, Mavinkurve M, Mendillo M, Watkins M, Gamieldien H, et al. Novel application of a whole blood intracellular cytokine detection assay to quantitate specific T-cell frequency in field studies. Journal of immunological methods. 2004;291(1-2):185–95.

30. Netea MG, Quintin J, van der Meer JW. Trained immunity: a memory for innate host defense. Cell Host Microbe. 2011;9(5):355–61.

31. Netea MG, Domínguez-Andrés J, Barreiro LB, Chavakis T, Divangahi M, Fuchs E, et al. Defining trained immunity and its role in health and disease. Nature Reviews Immunology. 2020;20(6):375–88.

32. Martinez FO, Gordon S. The M1 and M2 paradigm of macrophage activation: time for reassessment. F1000prime reports. 2014;6.

33. Jamil B, Shahid F, Hasan Z, Nasir N, Razzaki T, Dawood G, et al. Interferonγ/IL10 ratio defines the disease severity in pulmonary and extra pulmonary tuberculosis. Tuberculosis. 2007;87(4):279–87.

34. Zhang C, Yin J, Zheng J, Xiao J, Hu J, Su Y, et al. EZH2 identifies the precursors of human natural killer cells with trained immunity. Cancer biology & medicine. 2021;18(4):1021.

35. Oleszycka E, McCluskey S, Sharp FA, Muñoz-Wolf N, Hams E, Gorman AL, et al. The vaccine adjuvant alum promotes IL-10 production that suppresses Th1 responses. European Journal of Immunology. 2018;48(4):705–15.

36. Mezouar S, Diarra I, Roudier J, Desnues B, Mege J-L. Tumor necrosis factor-alpha antagonist interferes with the formation of granulomatous multinucleated giant cells: new insights into Mycobacterium tuberculosis infection. Frontiers in Immunology. 2019;10:1947.

37. Kleinnijenhuis J, Quintin J, Preijers F, Joosten LA, Jacobs C, Xavier RJ, et al. BCG-induced trained immunity in NK cells: role for non-specific protection to infection. Clinical immunology. 2014;155(2):213–9.

38. Fujihara M, Muroi M, Tanamoto K-i, Suzuki T, Azuma H, Ikeda H. Molecular mechanisms of macrophage activation and deactivation by lipopolysaccharide: roles of the receptor complex. Pharmacology & therapeutics. 2003;100(2):171–94.

39. Moorlag S, Arts R, Van Crevel R, Netea M. Non-specific effects of BCG vaccine on viral infections. Clinical microbiology and infection. 2019;25(12):1473–8.

40. Rousset F, Garcia E, Defrance T, Peronne C, Vezzio N, Hsu D-H, et al. Interleukin 10 is a potent growth and differentiation factor for activated human B lymphocytes. Proceedings of the National Academy of Sciences. 1992;89(5):1890–3.

41. Obukhanych TV, Nussenzweig MC. T-independent type II immune responses generate memory B cells. J Exp Med. 2006;203(2):305–10.

42. Young D, Roman E, Moreno C, O’Brien R, Born W. Molecular chaperones and the immune response. Philosophical Transactions of the Royal Society of London Series B: Biological Sciences. 1993;339(1289):363–8.

43. Heck TG, Ludwig MS, Frizzo MN, Rasia-Filho AA, Homem de Bittencourt Jr PI. Suppressed anti-inflammatory heat shock response in high-risk COVID-19 patients: lessons from basic research (inclusive bats), light on conceivable therapies. Clinical Science. 2020;134(15):1991–2017.

