## supplemental tables for "BCG activation of trained immunity is associated with induction of cross reactive COVID-19 antibodies in a BCG vaccinated population"

Karachi 74800, Pakistan

**Supplementary Material: Figures**

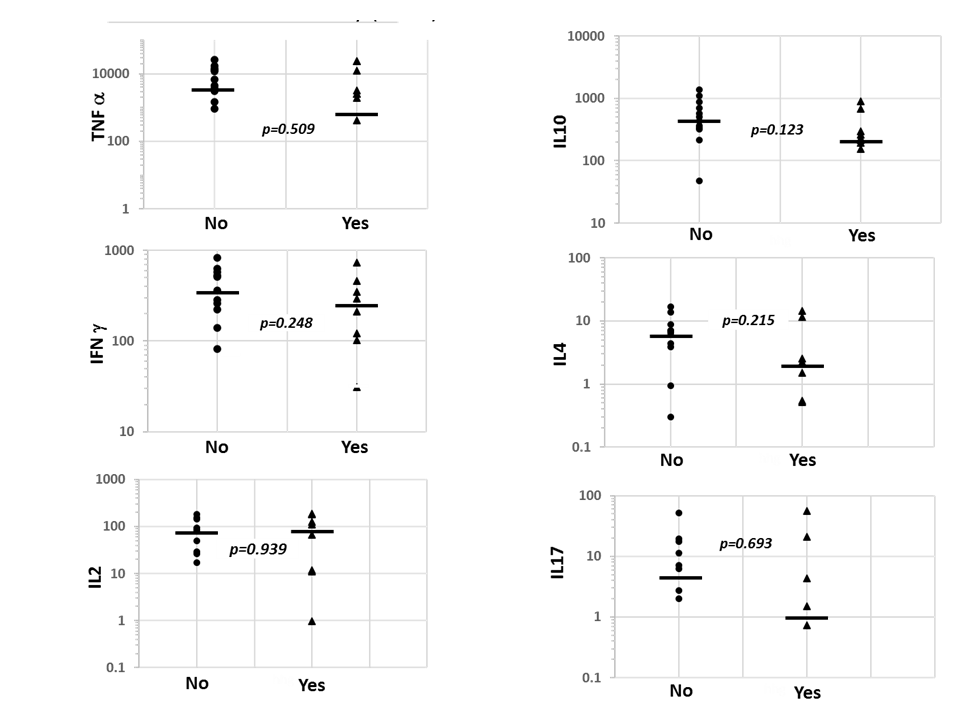

**Supplementary Fig. S1. Comparison of cytokine secretion in relation to Bacille Calmette-Guerin (BCG) status in whole blood assay (WBA)**

Comparison of cytokine secretion in BCG scar positive (n=8) and scar negative (n=12) Scatter plot shows individual responses of cytokines in both groups. Results are shown as individual data points for stimulated cytokines in whole blood assay (WBA). Mann-Whitney U test was applied to compare Scar positive and negative groups. The solid bars compare the median responses in the two groups.

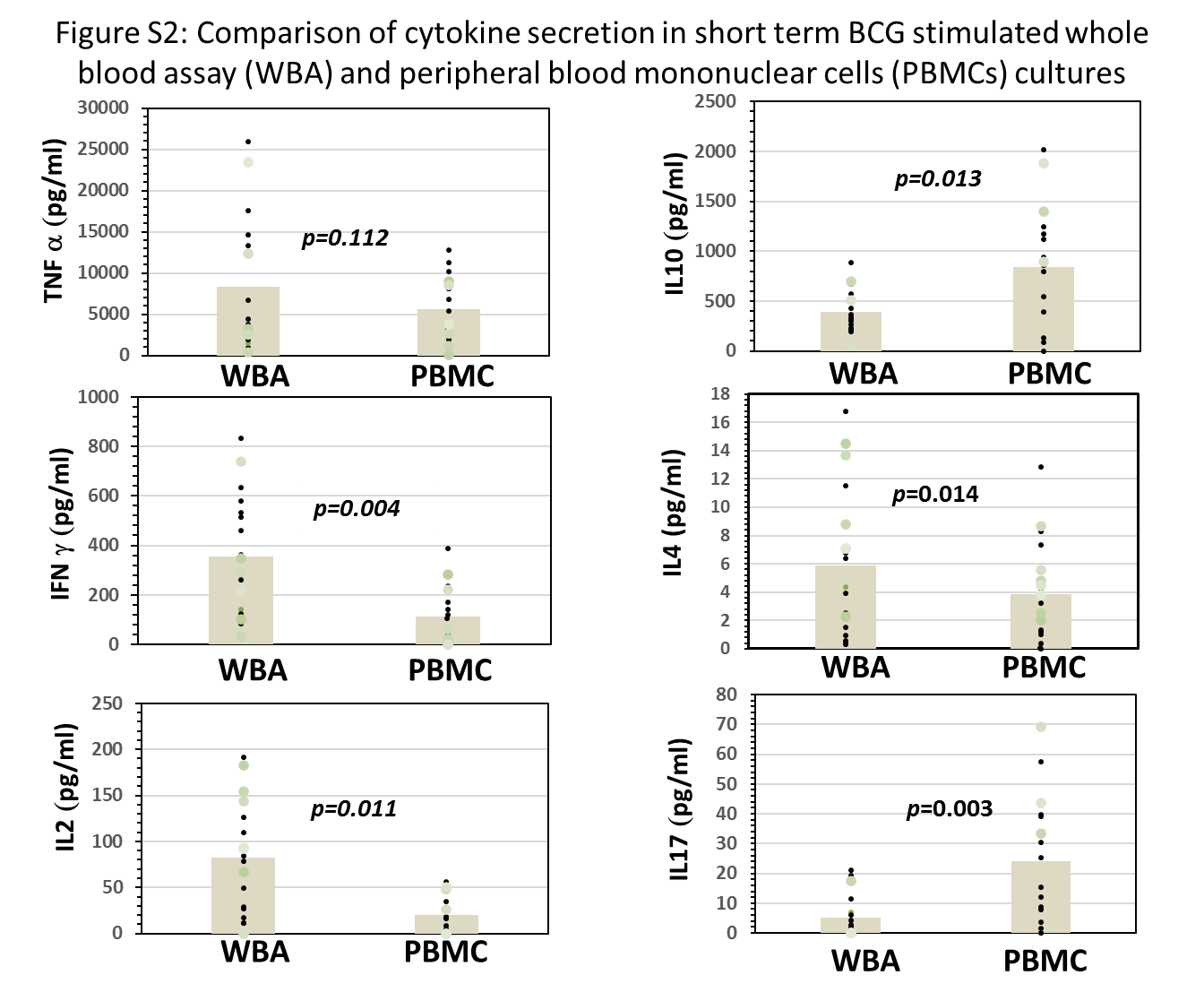

**Supplementary Fig. S2. Comparison of cytokine secretion in short term BCG stimulated whole blood assay (WBA) and peripheral blood mononuclear cells (PBMCs) cultures**

Comparison of cytokine secretion in short term WBA (12 hrs.) and PBMC (12 hrs.) BCG stimulated cultures (n=20)**.** Whole blood was diluted 1/2 in complete RPMI media and stimulated with 60ul of BCG (2-8 x10^6 CFU/vial), to obtain an MOI of 1.2. For PBMC a concentration of ~2 million cells were used for the stimulation of cells. The results are shown as log values after deducting spontaneous secretion (pg/ml). The shaded bars indicate the mean levels. The Wilcoxon Sign Rank test was applied for the significant difference in the cytokine level. p<0.05 was considered significant.

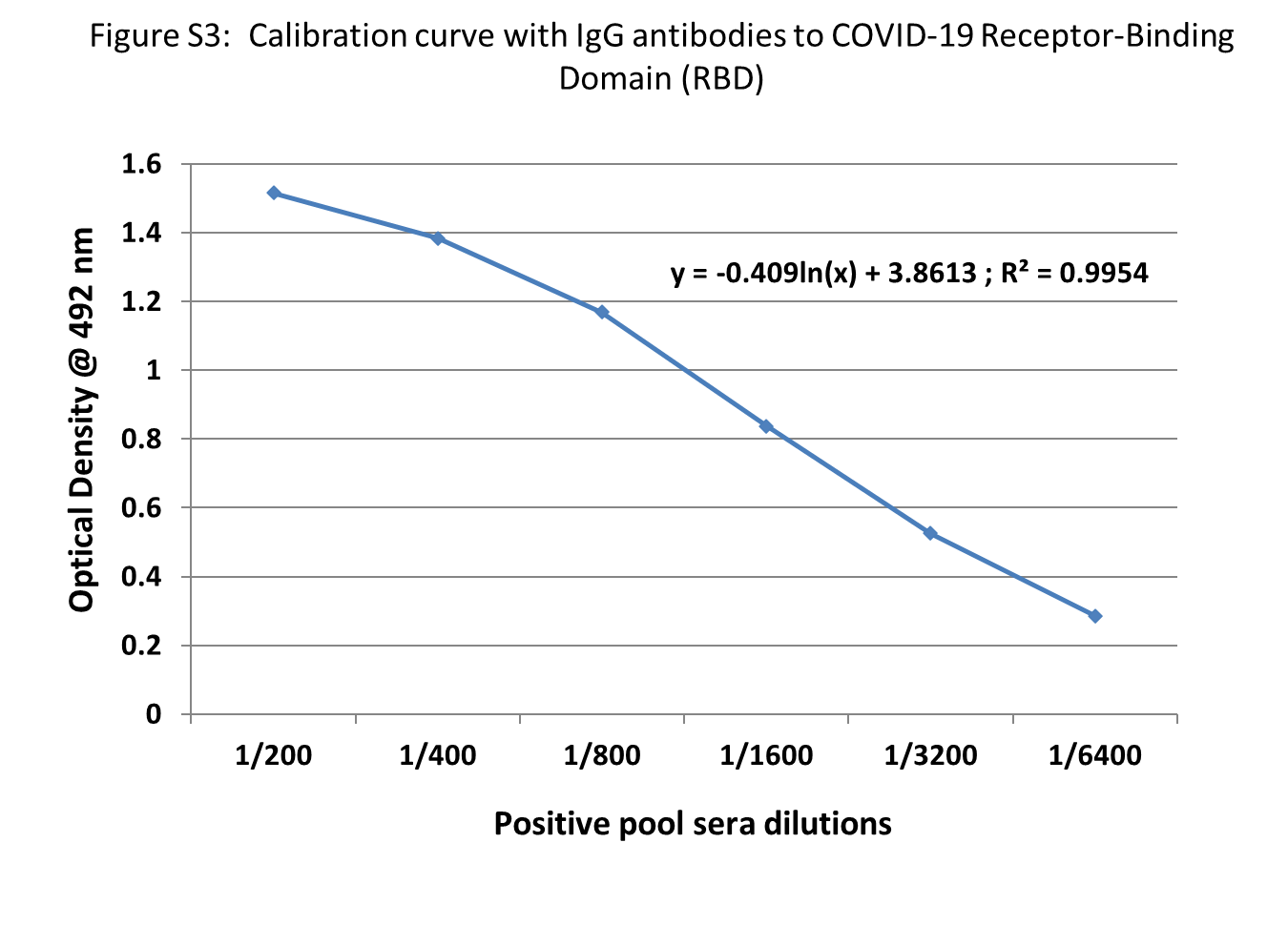

**Supplementary Fig. S3. Calibration curve with IgG antibodies to COVID-19 Receptor binding domain (RBD)**

A positive pool for IgG anti-RBD antibodies was diluted to determine endpoint titer. Endpoint titer was taken as one unit of antibody activity.

**Supplementary Material Tables**

**Supplementary Table S1: Cells and inducers of cytokines in the Innate arm of the immune system**

| Subset of Immune cells | Inducers | Cytokines secreted |
| --- | --- | --- |
| M1 (inos) ^a^ | LPS, TNF | TNFα, IFNγ |
| M2A (arg) ^a^ | IL4 | IL10 |
| M2B (inos) ^a^ | LPS | IL10, TNFα |
| M2C (arg) ^a^ | IL10 | IL10 |
| γδ T cells (BTN3A2) ^b^ | BCG stress proteins LPS | IL4, IL17 |
| NK cells (GZMA) ^c^ | LPS, BCG | IL2, IFNγ, TNFα, IL10 |

a (1, 2), b (3, 4), c (5)

**Note: Cytokine and inducers analyzed in the current study**

**Supplementary Table S2: Age-wise (minimum to maximum) BCG scar status, Mantoux Test (MT) readings, and gender distribution of study participants**

| S # | Participant ID | Age (years) | Gender | BCG Scar | MT reading (mm) |
| --- | --- | --- | --- | --- | --- |
| 1 | SMG-20 | 16 | Female | Yes | 0 |
| 2 | SMG-21 | 16 | Female | No | 7 |
| 3 | SMG-12 | 17 | Male | Yes | 0 |
| 4 | SMG-13 | 17 | Male | No | 2 |
| 5 | SMG-22 | 20 | Male | No | 0 |
| 6 | SMG-23 | 20 | Male | Yes | 0 |
| 7 | SMG-10 | 23 | Female | Yes | 0 |
| 8 | SMG-11 | 23 | Female | No | 0 |
| 9 | SMG-24 | 27 | Male | Yes | 0 |
| 10 | SMG-25 | 27 | Male | No | 0 |
| 11 | SMG-14 | 28 | Female | No | 0 |
| 12 | SMG-15 | 28 | Female | Yes | 4 |
| 13 | SMG-18 | 28 | Female | Yes | 0 |
| 14 | SMG-19 | 28 | Female | No | 0 |
| 15 | SMG-26 | 34 | Male | No | 0 |
| 16 | SMG-27 | 34 | Male | No | 0 |
| 17 | SMG-16 | 35 | Female | Yes | 0 |
| 18 | SMG-17 | 35 | Female | No | 0 |
| 19 | SMG-28 | 40 | Male | No | 0 |
| 20 | SMG-29 | 40 | Male | No | 0 |

**Supplementary Table S3: Primers for detecting gene expression in BCG activated cells of the innate immune system**

| S No | Target Gene* | Primer Sequence (5’ 3’) |
| --- | --- | --- |
| 1 | GZMA | Forward: TTT CTG GCA TCC TCT CTC TCA |
|  |  | Reverse: GGG TCA TAG CAT GGA TAG GG |
| 2 | BTN3A2 | Forward: AAG ACA GCC AGC ATT TCC AT |
|  |  | Reverse: GAG AAG CAG CAG CAA GAT AGG |
| 3 | TNFα | Forward: AGC CCA TGT TGT AGC AAA CC |
|  |  | Reverse: TGA GGT ACA GGC CCT CTG AT |
| 4 | 36B4 | Forward: TCC TCT CAC CAG GTG TCG TC |
|  |  | Reverse: CTG TCT TCC CTG GGC ATC AC |

*****GZMA (Granzyme A); marker for natural killer (NK) cells. BTN (Butyrophilin); marker for gamma-delta (γδ) T cells. TNFα (Tumor Necrosis Factor); marker for M1 cells

**Supplementary Table S4. Comparison of spontaneous secretion of cytokines in whole blood assay (WBA) and peripheral blood mononuclear cells (PBMCs) cultures**

| Cytokines (pg/ml) * | WBA (n=20) | PBMC (n=20) | *p-*value ^ɸ^ |
| --- | --- | --- | --- |
| IL-2 (mean ± SE) | 3.2 ± 2.8 | 0.46 ± 0.20 | 0.500 |
| IL-4 (mean ± SE) | 0.09 ± 0.06 | 0.04 ± 0.02 | 0.859 |
| IL-17(mean ± SE) | 0.03 ± 0.03 | 0.10 ± 0.10 | 0.317 |
| IL-10 (mean ± SE) | 12.1 ± 9.1 | 15.3 ± 10.6 | 0.017 |
| IFNγ (mean ± SE) | 12.7 ± 7.7 | 8.7 ± 5.4 | 0.715 |
| TNFα (mean ± SE) | 69.2 ± 67.1 | 50.0 ± 23.9 | 0.009 |

***** IL, interleukin; SE, standard error; IFNγ, interferon gamma; TNFα, tumor necrosis factor alpha

^ɸ^ *p* value <0.05 considered as significant value

**Supplementary Table S4.** Cytokine mean levels and standard error around the means are given. Cytokines were assessed by the Luminex assay system. Wilcoxon sign rank test was applied for the significant difference in spontaneous secretion of cytokines in WBA (12hrs) and PBMC (12hrs) cultures. Mann-Whitney U test was carried out to determine the significance of differences in the two assays.

**Supplementary Table S5. The magnitude of BCG recall responses in Trained Immunity (TI)**

| Cytokines* | WBA ratio (stimulated / spontaneous) | PBMCs ratio (stimulated / spontaneous) |
| --- | --- | --- |
| TNFα | 132 (9148 / 69) | 113 (5640 / 50) |
| IFNγ | 27 (355 / 13) | 132 (132 / 1) |
| IL-10 | 40 (488 / 12) | 56 (842 / 15) |
| IL-2 | 27 (83 / 3) | 24 (24/ 1) |
| IL-4 | 6 (6 / 1) | 4 (4 / 1) |
| IL-17 | 10 (10 / 1) | 24 (24 / 1) |

* TNFα, tumor necrosis factor alpha; IFNγ, interferon gamma; IL, interleukin

**Supplementary Table S5.** BCG recall responses in whole blood assay (WBA) and peripheral blood mononuclear cells (PBMCs) were determined by the ratio of stimulated cytokines levels divided by spontaneous cytokines levels. M1 derived cytokines (TNFα, IFNγ); M2 derived (IL10); NK cells (IL2), gamma-delta (γδ) T cells (IL4, IL17), (refer to supplementary Table S1). Cytokines <1.0 pg/ml were rounded off to 1.0

**Supplementary Table S6. Correlation between cytokines secretions in response to Bacille Calmette-Guerin in Whole Blood Assay (WBA) culture**

| Cytokines Correlation Coefficient | IL2 BCG_12hrs WBA | IL4 BCG_12hrs WBA | IL10 BCG_12hrs WBA | IL17 BCG_12hrs WBA | IFNγ BCG_12hrs WBA | TNFα BCG_12hrs WBA |
| --- | --- | --- | --- | --- | --- | --- |
| IL2 BCG_12hrs WBA | 1.000 | .642^**^ | .277 | .676^**^ | .674^**^ | .642^**^ |
| IL4 BCG_12hrs WBA | .642^**^ | 1.000 | .515^*^ | .647^**^ | .838^**^ | .860^**^ |
| IL10 BCG_12hrs WBA | .277 | .515^*^ | 1.000 | .361 | .367 | .362 |
| IL17 BCG_12hrs WBA | .676^**^ | .647^**^ | .361 | 1.000 | .711^**^ | .528^*^ |
| IFNγ BCG_12hrs WBA | .674^**^ | .838^**^ | .367 | .711^**^ | 1.000 | .759^**^ |
| TNFα BCG_12hrs WBA | .642^**^ | .860^**^ | .362 | .528^*^ | .759^**^ | 1.000 |

** Correlation is significant at the 0.01 level (2-tailed)

* Correlation is significant at the 0.05 level (2-tailed)

**Supplementary Table S7. Correlation between cytokine secretions in response to Bacille Calmette-Guerin in Peripheral Blood Mononuclear Cells (PBMCs) culture**

| Cytokines Correlation Coefficient | IL2 BCG_48hrs PBMC | IL4 BCG_48hrs PBMC | IL10 BCG_48hrs PBMC | IL17 BCG_48hrs PBMC | IFNγ BCG_48hrs PBMC | TNFα BCG_48hrs PBMC |
| --- | --- | --- | --- | --- | --- | --- |
| IL2 BCG_48hrs PBMC 1.000 | | .700** | .599* | .744** | .717** | .696** |
| IL10 BCG_48hrs PBMC | .599* | .570* | 1.000 | .681** | .455 | .541* |
| IL17 BCG_48hrs PBMC | .744** | .941** | .681** | 1.000 | .862** | .911** |
| IFNγ BCG_48hrs PBMC | .717** | .923** | .455 | .862** | 1.000 | .961** |
| TNFα BCG_48hrs PBMC | .696** | .950** | .541* | .911** | .961** | 1.000 |
| IL4 BCG_48hrs PBMC | .700** | 1.000 | .570* | .941** | .923** | .950** |

** Correlation is significant at the 0.01 level (2-tailed)

* Correlation is significant at the 0.05 level (2-tailed)

**Supplementary Table S8. Spearman Rank correlation between BCG and LPS stimulated cytokines secreted in whole blood assay (WBA) culture**

| BCG vs LPS | IL2 | IL4 | IL10 | IL17 | IFNγ | TNFα |
| --- | --- | --- | --- | --- | --- | --- |
| rho | 0.562* | 0.611** | -0.033 | 0.273 | 0.464* | 0.670** |
| *p* | 0.015 | 0.005 | 0.896 | 0.258 | 0.045 | 0.002 |
| n | 18 | 19 | 18 | 19 | 19 | 19 |

** Correlation is significant at the 0.01 level (2-tailed)

* Correlation is significant at the 0.05 level (2-tailed)

*p-*value <0.05 considered as significant value

**References**

1.	Foey AD. Macrophages—masters of immune activation, suppression and deviation. Immune response activation. 2014;276.

2.	Martinez FO, Gordon S. The M1 and M2 paradigm of macrophage activation: time for reassessment. F1000prime reports. 2014;6.

3.	Hoft DF, Brown RM, Roodman ST. Bacille Calmette-Guérin vaccination enhances human γδ T cell responsiveness to mycobacteria suggestive of a memory-like phenotype. The Journal of Immunology. 1998;161(2):1045-54.

4.	Wo J, Zhang F, Li Z, Sun C, Zhang W, Sun G. The role of gamma-delta T cells in diseases of the central nervous system. Frontiers in immunology. 2020;11:580304.

5.	Kleinnijenhuis J, Quintin J, Preijers F, Joosten LA, Jacobs C, Xavier RJ, et al. BCG-induced trained immunity in NK cells: role for non-specific protection to infection. Clinical immunology. 2014;155(2):213-9.
